# Validation of the Malawi Developmental Assessment Tool for children in the Dominican Republic

**DOI:** 10.1101/497347

**Authors:** Laura V. Sánchez-Vincitore, Paul Schaettle, Arachu Castro

## Abstract

**Background:** This study initiated the validation process of a translated and adapted version of the Malawi Developmental Assessment Tool (MDAT) for children in the Dominican Republic (DR). Like Malawi before the development of the MDAT, the DR did not have early childhood development (ECD) tools explicitly designed for low-resource areas that are also valid assessments of child development. We chose MDAT because it underwent a rigorous validation process and retained measurements of test items that were culturally adaptable from the Denver Developmental Screening Test II. We aimed to test the internal consistency and inter-rater reliability of the psychometric properties of the MDAT in children under the age of two years living in low-income neighborhoods in Santo Domingo in 2017.

**Methods and Findings:** Forty-two children from 2 to 24 months of age (mean = 11.26, SD = 6.37, boys = 22, girls = 20) and their corresponding caregiver participated in the study. We conducted a cross-sectional, pre-experimental study. The primary outcome measure was an index of ECD, as assessed by the Dominican adaptation of the MDAT. The tool evaluates children in four domains: social, fine motor, language, and gross motor. To determine internal consistency, we obtained Cronbach’s alpha for each sub-scale. The results ranged from 0.89 to 0.94, indicating good consistency. Second, to test the interrater reliability, we conducted a Kendall’s Taub test of independence for both the general scale and each sub-scale. Significant rτ scores ranged from .923 to .966, indicating appropriate interrater reliability. Third, we correlated the age variable with each subscale to determine if the development scale followed a progression of abilities that are expected to increase with maturation. The age variable correlated positively with all the subscales (social r=.887, p < .001; fine motor r = .799, p < .001; language r = .834, p < .001; gross motor r = .805, p < .001), indicating that the older the child, the better scores in the development measurements, as expected. There were no adverse events. This study, however, has multiple limitations. We did not gather information about socioeconomic position, which is an important variable when assessing child development; however, all participants lived in a low-income neighborhood. Given that this is the first ECD tool specific to the Dominican Republic, norm-referenced scores for the Dominican population do not yet exist. This study sample size is insufficient to make inferences about the national population.

**Conclusions:** This study represents the first attempt to obtain a valid tool to screen for development milestones in children living in poverty in the DR. More research is needed to refine the instrument. The availability of the tool will enable impact evaluations of ECD intervention programs and the development of evidence-based public policies in the DR.

## Introduction

Developmental assessment or screening tools provide a standardized method of assessing a child’s neurological and musculoskeletal growth through the observation of the child’s performance of age and culturally-specific activities (1). The child is observed performing a set of tasks associated with specific interrelated domains and evaluated based on direct structured observations of the expected behavior, caregiver reports, or unstructured observation from evaluators (2). As the assessment progresses, the child engages in activities of increasing difficulty (2).

There are numerous benefits associated with the availability and use of developmental screening tools. At the individual level, these screening tools help determine if a child is on track in his or her development, identify interventions to compensate for any eventual delay, and implement early interventions that help improve their health and educational outcomes (3). At the program level, developmental screening tools are used as baseline and outcome variables in impact evaluations to help determine a program’s effectiveness (4). At the public policy level, the use of screening tools helps guide the development of evidence-based health and education policies (5).

Several tools have been created to measure early childhood development (ECD) in a range of domains, standardized with large representative samples in places that have health data readily available, piloted, and validated. These data-backed assessments of the tools’ ability to assist health professionals in the measurement of ECD make them appropriate resources for assessing different aspects of child development in those locations (6). Despite the availability of these tools and their translation into a variety of languages, they may not necessarily be adequate to measure ECD in cultural and socioeconomic contexts for which the instruments were not specifically created. For example, a study in Chile adapted the Bayley-III developmental tool and validated it with a sample of children from higher socioeconomic position families, which was “representative of the private medical center where the study was conducted” (7). This shows that while the adapted screening tool was valid for that specific context, it was not necessarily applicable to lower socioeconomic position participants regardless of their shared geographic location and language. For this reason, it is essential to ensure that development tools are designed with the input of participating communities and validated with a sample representative of the specific population in which it will be used.

Children’s development depends on multiple factors, including childrearing practices that are culture-specific. Therefore, using development tools without validating them in the cultural and socioeconomic context where they will be used can lead to an under- or over-estimation of ECD (8). Some experiences exist across the world of contextualized ECD screening tools for specific populations in India, Pakistan, and Zambia (9), Malawi (10), Sri Lanka (11), Cambodia (12), and Aboriginal Australia (13). These tools were designed and validated with as many culture-free items as deemed possible, but also with items that account for specific population characteristics of environments that are frequently not represented by the most commonly used developmental screening tools.

In addition to having a more culturally relevant measurement to assess ECD, it is necessary for these screening tools to be accessible for projects, programs, and research at the national level. The accessibility guarantees the constant use of the instrument and the standardization of ECD measurement across projects. Therefore, commercial ECD screening tools used to measure development or to screen for developmental delay in children are expensive and are used mostly in clinical settings (14). Tools that can help health professionals in these areas identify at-risk children for developmental delays and assess if they are developing according to their age need to be available at low or no cost to the provider to maximize their use (6).

The purpose of our study was to test an ECD tool that could be used in the Dominican Republic (DR) at the community level in a resource-poor setting and no cost. The DR faces multiple challenges in educational attainment, as reflected by international educational reports, which show that Dominican students have the lowest scores from a subset of fifteen Latin American countries in reading, writing, and math in third and sixth grade (15). An early literacy national study conducted in 2015 showed that second graders had still not acquired basic literacy skills (16), partly due to low oral comprehension—a skill that the education system implicitly assumes the child has acquired before entering formal educational settings (17). On the other hand, no ECD testing tools have been developed specifically for the Dominican context, as the only ones that are used are available in private clinics, such as the Developmental Profile 3 (18) and the Denver Developmental Screening Test II (DDST-II) (19).

In our study, we aimed to test the internal consistency and inter-rater reliability of the psychometric properties of the Malawi Development Assessment Tool (MDAT) (10) in a group of children under the age of two years in the Dominican Republic. The MDAT is an ECD screening tool that focuses on a continuum of skills from four different domains—gross motor, fine motor, language, and social, with the purpose of identifying children with severe disabilities. Like Malawi before the development of the MDAT, the Dominican Republic does not have ECD tools designed specifically for low-resource areas in the country that are also valid assessments of child development. After reviewing a variety of ECD screening tools, we chose the MDAT because it was developed for children ages 0 to 5 years, underwent a rigorous validation process informed by Malawian health workers and pediatricians, and retained measurements of test items that were culturally adaptable (6, 10) from the Denver Developmental Screening Test II (DDST-II) (19)—which is one of the most used instruments to assess child development in a short amount of time and that can be used by “anyone who works well with children and meticulously follows directions for administration” (6). These qualities are ideal for use in low-resource environments where many children must be assessed quickly and highly-trained health care workers are not available.

## Methods and materials

### Participants

Forty-two children from 2 to 24 months of age (mean = 11.26, SD = 6.37, boys = 22, girls = 20) and their corresponding caregiver—their mother in all the cases—participated in the study. We recruited study participants in Los Guandules and Guachupita, two neighborhoods of high economic deprivation in the Santo Domingo metropolitan area, the capital city of the DR. Inclusion criteria for the study were being a child from 0 to 24 months of age with a parent or guardian aged 18 years or older who understood Spanish—regardless of whether their first language was Spanish or Haitian Kreyol. Since our goal was to determine the validity of a tool that measures typical ECD, we excluded children with diagnosed developmental disabilities.

Volunteers from the Pastoral Materno Infantil (PMI), a Jesuit organization that promotes maternal and child health among low-income families throughout the Dominican Republic through trained community mobilizers who live in the community, recruited participants via convenience sampling by a phone call from the pool of PMI beneficiaries. Once the community mobilizers had identified a group of participants interested in the study, they gathered them and brought them to the evaluation setting. The Institutional Review Boards from the Universidad Iberoamericana (UNIBE) in Santo Domingo and Tulane University approved the study. We obtained oral and written consent from the child’s caregiver before data collection.

### Instruments

#### Sociodemographic interview

The interview consisted of three parts to assess participants’ background and their home environment: (a) information related to the child, including prenatal and perinatal history, access to stimulating materials such as books and toys, and interaction with other people such as singing, speaking, and storytelling; (b) information about the primary caregiver, including level of education and the relationship with the child; (c) general nutritional indicators such as the source of household’s water for cooking, cleaning, and drinking.

#### Malawi Development Assessment Tool – Dominican version

At our request, the MDAT team provided us with materials to assist in our adaptation with thorough documentation of the process they underwent to create and validate the tool. We translated the MDAT into Spanish from English by first directly translating the MDAT and then reviewing this version with community volunteers from PMI. As part of the assessment of the translation, we adapted the tasks of the original MDAT to the Dominican context by accounting for different availability of materials and participants’ familiarity with certain activities. We named this new version of the test MDAT-DR. The appropriateness of the choice of words used and tasks involved in the MDAT-DR were informed by discussions with staff and volunteers from PMI.

The MDAT-DR consists of four subtests that assess development in four different domains: social, fine motor, language, and gross motor. Each subtest contains a list of 34 items of behaviors that progress in complexity. Each item is scored with categorical answers 0, 1, or 2. A score of 1 is given if the evaluator observed the behavior, a 2 if the caregiver reported that the child performs the task, and 0 for behaviors that were neither observed by the evaluator nor reported by the caregiver as having been performed. The child’s age determined the starting point of each domain. Each item was tested and scored as “pass observed,” “pass reported,” or “fail.” We administered the items sequentially,. When the child failed to complete six tasks in a row, the evaluator moved on to the next subtest.

### Procedure

Data collection took place throughout eight days in February 2017 at Centro Bonó— another Jesuit center in the same sector of Santo Domingo. A group of nine evaluators conducted the assessments in three separate rooms; two evaluators assessed each child and each of them provided their own set of scores. These evaluators were clinical psychology undergraduate students from UNIBE who had already completed research and ECD measurement courses. The local principal investigator (PI), a neuroscientist, provided them with a 4-hour training on the study protocol, participant protection, and childhood development, and supervised them when interacting with participants to ensure participant safety and study integrity.

First, the evaluators conducted the sociodemographic interview with each caregiver using a structured multiple-choice questionnaire that took approximately 10 minutes. Upon completion of the interview, the evaluators administered the MDAT-DR under the supervision of the local PI. Once the evaluators completed data collection, the data entry team consisting of UNIBE undergraduate psychology students inputted the data, which were reviewed by the local PI. We converted the data to a binary scale, with “fail” coded as 0 and both “pass reported” and “pass observed” coded as 1. By numerically adding the “pass” responses, each child received a score from 0 to 34 on a continuum for each subscale. We analyzed the scores for internal consistency and inter-rater reliability.

## Results

### Sociodemographic information

The age of the 42 children who participated in the study ranged between 2 and 24 months, as shown in Table 1.

**Table 1:** Age and sex distribution of children participants, Santo Domingo, 2017

| Age (in months) | Female | Male |
| --- | --- | --- |
| 2 | 0 | 2 |
| 3 | 3 | 1 |
| 4 | 0 | 1 |
| 5 | 1 | 1 |
| 6 | 1 | 1 |
| 7 | 1 | 1 |
| 8 | 3 | 3 |
| 9 | 1 | 2 |
| 12 | 1 | 2 |
| 14 | 1 | 0 |
| 15 | 1 | 0 |
| 16 | 2 | 3 |
| 17 | 1 | 2 |
| 18 | 1 | 0 |
| 19 | 0 | 2 |
| 20 | 1 | 0 |
| 22 | 1 | 0 |
| 24 | 1 | 1 |
| Total | 20 | 22 |

According to their caregivers, 23.8% of the children were born with low birth weight, and 16.7% were born prematurely. When asked who was regularly in charge of caring for their children, most reported that the main caregiver was the mother, followed by mother and father, and the mother and grandmother (see Table 2). The results show that 23.8% of the children’s mothers had elementary education level, 64.3% had secondary school education level, and 11.9% attended college. Forty-one caregivers spoke Spanish as a first language, and one caregiver spoke Haitian Kreyol as a first language. All of the households used bottled water to drink; to cook, 21 used tap water, 19 used bottled water, and 2 used water bought from a delivery truck; for cleaning and bathing, all the households used tap water, and one used water from the *camioncito* [a water delivery truck].

**Table 2:** Frequency of primary caregiver in a sample of 42 children ages from 2 months to 2 years, Santo Domingo, 2017

| **Primary caregiver** |  |
| --- | --- |
| Mother | 61.9% |
| Mother and father | 16.7% |
| Mother and grandmother | 11.9% |
| Grandmother | 4.8% |
| Sister | 2.4% |
| Mother and other | 2.4% |
| Total | 100% |

Tables 3 and 4 depict the home background analysis, which includes access to stimulating materials and stimulating activities.

**Table 3:** Access to stimulating materials in children ages from 2 months to 2 years, Santo Domingo, 2017

| **Materials** | **N** | **Min** | **Max** | **Mean** | **SD** |
| --- | --- | --- | --- | --- | --- |
| Books at home | 41 | 0 | 2 | 0.3 | 0.656 |
| Toys at home | 37 | 1 | 20 | 6.3 | 4.235 |

Min = Minimum value; Max = Maximum value; M = Mean; SD = Standard deviation

**Table 4:** Access to stimulating activities in children ages from 2 months to 2 years, Santo Domingo, 2017

| **Activities** | **N** | **%** |
| --- | --- | --- |
| % of children whose caregivers read stories | 41 | 26.8% |
| % of children whose caregivers tell stories | 41 | 48.8% |
| % of children whose caregivers sing to them | 42 | 92.9% |
| % of children whose caregivers take them out for a walk | 42 | 97.6% |
| % of children whose caregivers play with them | 42 | 100.0% |
| % of children whose caregivers counts and name objects to them | 42 | 85.7% |

### MDAT psychometric properties

First, we analyzed the MDAT-DR’s internal consistency to determine the degree to which items within each sub-scale were correlated. We obtained Cronbach’s alpha for each subscale: social, gross motor, language, and fine motor. Table 5 contains general descriptive statistics of each sub-scale, in addition to internal consistency data. Cronbach’s alpha ranges from 0.89 to 0.94, indicating a good consistency (20).

**Table 5:** Social, fine motor, language, and gross motor scales in children ages from 2 months to 2 years, Santo Domingo, 2017

| **Sub-scale** | **N** | **Min** | **Max** | **M** | **SD** | **$\alpha$** |
| --- | --- | --- | --- | --- | --- | --- |
| Social | 41 | 5 | 28 | 14.3 | 5.32 | 0.90 |
| Fine motor | 42 | 1 | 22 | 13.1 | 5.25 | 0.91 |
| Language | 42 | 3 | 23 | 10.0 | 4.20 | 0.88 |
| Gross motor | 41 | 1 | 27 | 15.3 | 6.27 | 0.94 |

Min = Minimum number; Max = Maximum number; M = Mean, SD = Standard Deviation; $\alpha$ = Cronbach's alpha

Second, to test the inter-rater reliability to ensure that multiple observers would obtain similar scores, we conducted a Kendall’s Taub test of independence for the general scale, as well as for each sub-scale. Scores obtained by the first evaluator were not independent from scores obtained by the second evaluator in any of the test (social *r*_τ_ = 0.953, p < 0.001; fine motor *r*_τ_ = 0.923, p < 0.001; language *r*_τ_ = 0.966, p < 0.001; gross motor *r*_τ_ = 0.977, p < 0.001; total *r*_τ_ = 0. 954, p < 0.001). Our interpretation of these results is that the scale has appropriate inter-rater reliability.

### Correlations

We correlated the age variable with each subscale to determine if the development scale followed a progression of abilities that are expected to increase with maturation. The age variable correlated positively with all the subscales (social r=.887, p < .001; fine motor r = .799, p < .001; language r = .834, p < .001; gross motor r = .805, p < .001), indicating that the older the child, the better scores in the development measurements, as expected. See Figure 1 for a visual representation.

**Figure 1:**
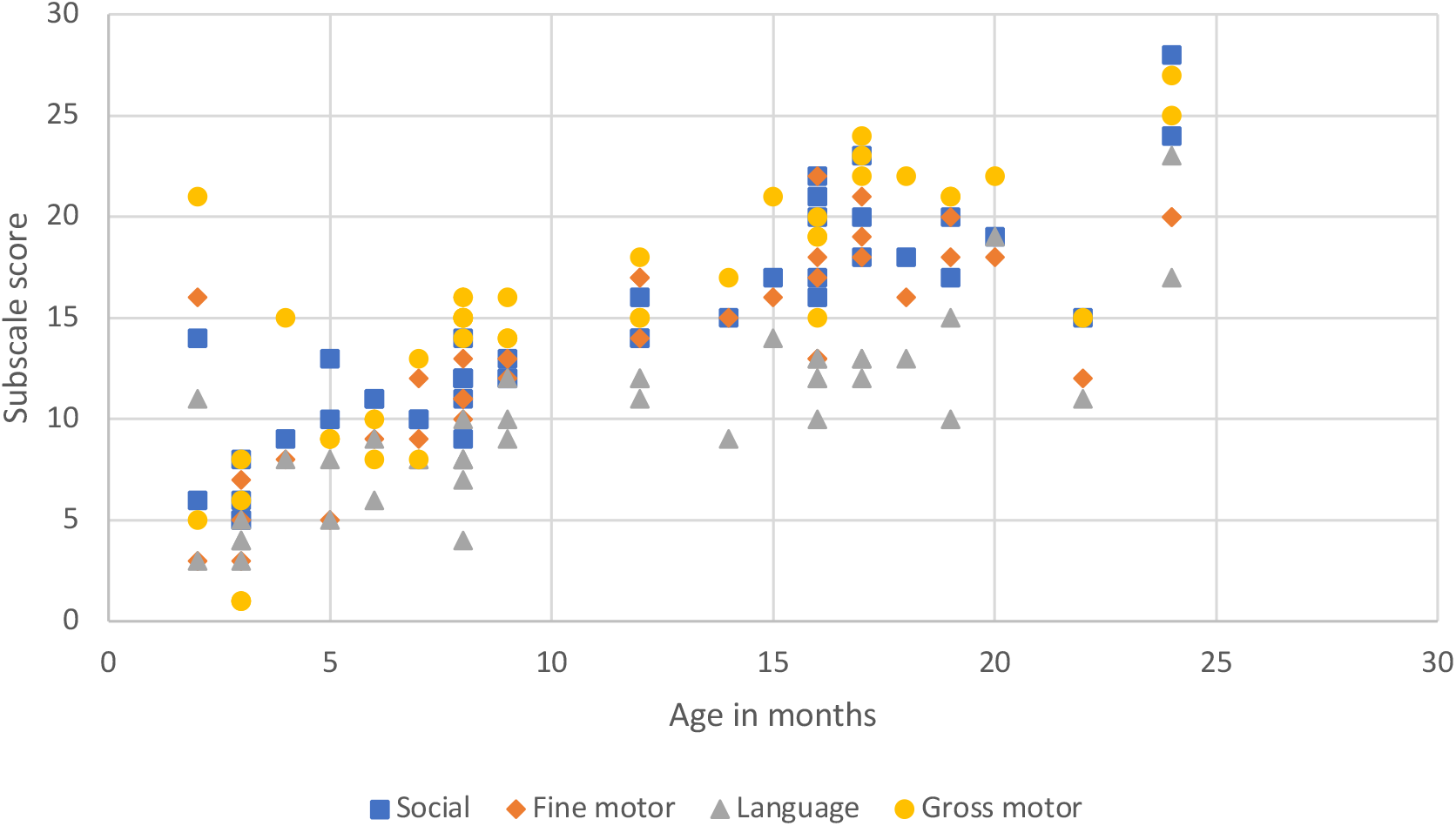
Age and developmental subscales correlation in 42 children ages from 2 months to 2 years, Santo Domingo, 2017

## Discussion

The primary objective of this study was to determine the feasibility of using an adapted version of the MDAT in a community context in vulnerable areas of Santo Domingo, Dominican Republic. We evaluated children that were between 2 months and 2 years of age, since items in developmental scales in such early stages are less culture-dependent and, therefore, require minimal adaptations. We obtained measurements of internal consistency of the instrument, as well as inter-rater reliability, while informally assessing the logistics and methodology of the study.

The instrument showed appropriate psychometric properties, including good internal consistency and good inter-rater reliability. This high index on internal consistency indicates a low probability of measurement errors from the design and content of the test itself. Good interrater reliability index indicates instrument stability across observers. By reducing error variance, threats to internal validity are reduced. As expected in any developmental scale that follows a path in child development, we confirmed a progression of scores as children were older.

Regarding logistics, one of the main strengths of this study was the affiliation with the Pastoral Materno Infantil. We chose PMI because they have a history of engaging the community and providing services that enable them to access health services. By partnering with PMI, we respected the way in which the community engages the health system. The evaluation setting was a space that participants already knew and visited regularly, and the parents trusted the community mobilizers who invited them to participate in the study. It would be interesting to explore the possibility of training the community mobilizers in the application of the screening tool, increasing the benefit of this project to the community and making this a sustainable community-engaged project.

This study, however, has multiple limitations. While there was no language requirement for participation by either participants or their caregivers, we observed that the child of the one caregiver with more limited Spanish abilities did not perform as well as the other children. This is because some questions were directly asked to parents, and if the parent did not understand the questions being asked, the child’s score could be affected negatively. Even though this was not common in this pilot study, for further studies in communities with immigrant populations, we recommend adding bilingual evaluators to the staff and additional translated materials to ensure appropriate representation of minority groups of languages.

The present study did not gather information about socioeconomic position, which is an important variable when assessing child development. The community coordinator and community mobilizers recruited participants from the same two neighborhoods, both of which include a large proportion of households under the local poverty line, but we did not take into account socioeconomic variability among the participants.

Because there are no available ECD tools specific to the Dominican Republic, there has been no developmental assessment on a national level. Therefore, we recommend the use of the MDAT-DR as an instrument to be used nation-wide to obtain norm-referenced scores for the Dominican population. The standardization of the scores would allow the use of the MDAT-DR for clinical and monitoring purposes at the community level. However, although developmental screening tools have the potential to infer about general development milestones, and probably detect children with significant impairments that require further testing, the use of screening tools may not be able to identify subtle developmental delays (2).

This study represents the first attempt to obtain a valid tool to screen for development milestones in children living in poverty in the Dominican Republic. More research is needed to refine the instrument, to have an available tool that is reliable and accessible to be used by health workers, and that could be used in future studies on factors that affect or enhance early childhood development. The availability of the tool will enable impact evaluations of early child development intervention programs and the development of evidence-based public policies on early childhood development in the Dominican Republic.

## Acknowledgments

We thank José Navarro and Altagracia Serrata from Pastoral Materno Infantil and Pablo Mella from Centro Bonó for facilitating the study. Special thanks to the evaluators, all of the students from the Neurocognition and Psychophysiology track at UNIBE: Analía Henríquez, Enmanuel Goico, Julia Tejeda, Ivanova Veras, Lorraine Ruiz, Aimée Castillo, Leslie Joaquín, Mélody Arias, and Maite Mejía, as well as to Patricia González, who contributed to data entering. Finally, we are thankful to Melissa Gladstone from the University of Liverpool, who shared the MDAT instrument and provided the MDAT training manual.

## References

1. Miller PJ, Goodnow JJ. Cultural practices: Toward an integration of culture and development. New Directions for Child and Adolescent Development. 1995;67:5–6.

2. Sabanathan S, Wills B, Gladstone M. Child development assessment tools in low-income and middle-income countries: how can we use them more appropriately? Arch Dis Child. 2015;100(5):482–8.

3. Rice CE, Naarden Braun KV, Kogan MD, Smith C, Kavanagh L, Strickland B, et al. Screening for developmental delays among young children--National Survey of Children’s Health, United States, 2007. MMWR Suppl. 2014;63(2):27–35.

4. Snow CE, Van Hemel SB, editors. Early Childhood Assessment: Why, What, and How. Washington, DC: The National Academy Press; 2008.

5. Wuermli AJ, Tubbs, C. C., Petersen, A. C., & Aber, J. L. Children and youth in low-and middle-income countries: Toward an integrated developmental intervention science. Child Development Perspectives. 2015;9(1):61–6.

6. Fernald L, Kariger P, Engle P, Raikes A. Examining early child development in low-income countries: A Toolkit for the Assessment of Children in the First Five Years of Life. Washington, DC: World Bank; 2009.

7. Schonhaut L, Armijo I, Schonstedt M, Alvarez J, Cordero M. Validity of the ages and stages questionnaires in term and preterm infants. Pediatrics. 2013;131(5):e1468–74.

8. Mendonça B, Sargent B, Fetters L. Cross-cultural validity of standardized motor development screening and assessment tools: a systematic review. Dev Med Child Neurol. 2016;58(12):1213–22.

9. Biasini FJ, De Jong D, Ryan S, Thorsten V, Bann C, Bellad R, et al. Development of a 12 month screener based on items from the Bayley II Scales of Infant Development for use in Low Middle Income countries. Early Hum Dev. 2015;91(4):253–8.

10. Gladstone M, Lancaster GA, Umar E, Nyirenda M, Kayira E, van den Broek NR, et al. The Malawi Developmental Assessment Tool (MDAT): the creation, validation, and reliability of a tool to assess child development in rural African settings. PLoS Med. 2010;7(5):e1000273.

11. Lokuketagoda BU, Thalagala N, Fonseka P, Tran T. Early Development Standards for Children Aged 2 to 12 Months in a Low-Income Setting. SAGE Open. 2016;6(4).

12. Ngoun C, Stoey LS, van’t Ende K, Kumar V. Creating a Cambodia-specific developmental milestone screening tool – a pilot study. Early Hum Dev. 2012;88(6):379–85.

13. Simpson S, D’Aprano A, Tayler C, Toon Khoo S, Highfold R. Validation of a culturally adapted developmental screening tool for Australian Aboriginal children: Early findings and next steps. Early Hum Dev. 2016;103:91–5.

14. Ringwalt S. Developmental Screening and Assessment Instruments with an Emphasis on Social and Emotional Development for Young Children Ages Birth through Five. Chapel Hill: The University of North Carolina, FPG Child Development Institute, National Early Childhood Technical Assistance Center; 2008.

15. UNESCO. Informe de Resultados TERCE, Tercer Estudio Regional Comparativo y Explicativo. Logros de aprendizaje [TERCE Results Report, Third Regional Comparative and Explanatory Study. Learning achievements]. Paris: UNESCO; 2016.

16. Mencía-Ripley A, Sánchez-Vincitore LV, Garrido LE, Aguasvivas-Manzano JA. Baseline report of USAID – Leer. Santo Domingo: USAID; 2016.

17. MINERD. Diseño Curricular Nivel Primario Primer Ciclo (1ro., 2do. y 3ro.) [Curricular Design Primary Level First Cycle (1st, 2nd, and 3rd)]. Santo Domingo: Ministerio de Educación de la República Dominicana; 2014.

18. Alpern GD. Developmental profile 3 (DP-3). Los Angeles: Western Psychological Services; 2007.

19. Frankenburg W, Dodds J, Archer P, Shapiro H, Bresnick B. Denver II technical manual. Denver: Denver Developmental Materials Inc.; 1990.

20. Kaplan RM, & Saccuzzo, D. Psychological Testing: Principles, Applications, and Issues, 6th edition. Belmont, California: Thomson Wadsworth; 2004.

